# Virological characteristics of the SARS-CoV-2 KP.3.1.1 variant

**DOI:** 10.1101/2024.07.16.603835

**Authors:** Yu Kaku, Keiya Uriu, Kaho Okumura, The Genotype to Phenotype Japan (G2P-Japan) Consortium, Jumpei Ito, Kei Sato

## Abstract

The SARS-CoV-2 JN.1 variant (BA.2.86.1.1), arising from BA.2.86.1 with spike protein (S) substitution S:L455S, outcompeted the previously predominant XBB lineages by the beginning of 2024. Subsequently, JN.1 subvariants including KP.2 (JN.1.11.1.2) and KP.3 (JN.1.11.1.3), which acquired additional S substitutions e.g., S:R346T, S:F456L, and S:Q493E, have emerged concurrently. Thereafter, JN.1 subvariants, such as LB.1 (JN.1.9.2.1), KP.2.3 (JN.1.11.1.2.3), and KP.3.1.1 (JN.1.11.1.3.1.1), which convergently acquired a deletion of Serine at the 31st position in S (S:S31del) in addition to the above substitutions, have emerged and spread as of June 2024. We recently reported the virological features of JN.1 subvariants including KP.2, KP.3, LB.1, and KP.2.3.2,3 Here, we investigated the virological properties of KP.3.1.1. First, we estimated the relative effective reproduction number (Re) of KP.3.1.1 using a Bayesian multinomial logistic model4 based on genome surveillance data from Spain, the USA, France, Canada, and the UK, where this variant has spread as of June 2024. In Spain, the Re of KP.3.1.1 is over 1.2-fold higher than that of JN.1 and even higher than those of KP.2, KP.3, LB.1, and KP.2.3. Additionally, the other countries under investigation herein show higher Re for KP.3.1.1. However, it must be noted there is the possibility of overestimation in these countries due to more limited KP.3.1.1 sequence numbers. These results suggest that KP.3.1.1 will spread worldwide along with other JN.1 sublineages. We then assessed the virological properties of KP.3.1.1 using pseudoviruses. The pseudovirus of KP.3.1.1 had significantly higher infectivity than that of KP.3. Neutralization of KP.3.1.1 was tested using i) convalescent sera after breakthrough infection (BTI) with XBB.1.5 or EG.5, ii) convalescent sera after the infection with HK.3 or JN.1, and iii) sera after monovalent XBB.1.5 vaccination. The 50% neutralization titer (NT50) against KP.3.1.1 was significantly lower than KP.3 (1.4–1.6-fold) in all four groups of convalescent sera tested. KP.3.1.1 also showed a 1.3-fold lower NT50 against XBB.1.5 vaccine sera than KP.3. Moreover, KP.3.1.1 showed stronger resistance with a 1.3-fold lower NT50 with statistical significances to the convalescent sera infected with EG.5 and HK.3 than KP.2.3. Altogether, KP.3.1.1 exhibited a higher Re, higher pseudovirus infectivity, and higher neutralization evasion than KP.3. These results align with our recent report that the JN.1 subvariants with S:S31del (e.g., KP.2.3 and LB.1) exhibited enhanced Re and immune evasion compared to the other JN.1 subvariants without S:S31del (e.g., JN.1, KP.2, and KP.3), highlighting the evolutionary significance of S:S31del in the JN.1 lineages.

## Text

The SARS-CoV-2 JN.1 variant (BA.2.86.1.1), arising from BA.2.86.1 with spike protein (S) substitution S:L455S, outcompeted the previously predominant XBB lineages by the beginning of 2024.^1^ Subsequently, JN.1 subvariants including KP.2 (JN.1.11.1.2) and KP.3 (JN.1.11.1.3), which acquired additional S substitutions e.g., S:R346T, S:F456L, and S:Q493E, have emerged concurrently (**Figure 1A**).^2^ Thereafter, JN.1 subvariants, such as LB.1 (JN.1.9.2.1), KP.2.3 (JN.1.11.1.2.3), and KP.3.1.1 (JN.1.11.1.3.1.1), which convergently acquired a deletion of Serine at the 31st position in S (S:S31del) in addition to the above substitutions, have emerged and spread as of June 2024 (**Figure 1A**). We recently reported the virological features of JN.1 subvariants including KP.2, KP.3, LB.1, and KP.2.3.^2,3^ Here, we investigated the virological properties of KP.3.1.1.

**Figure 1.**
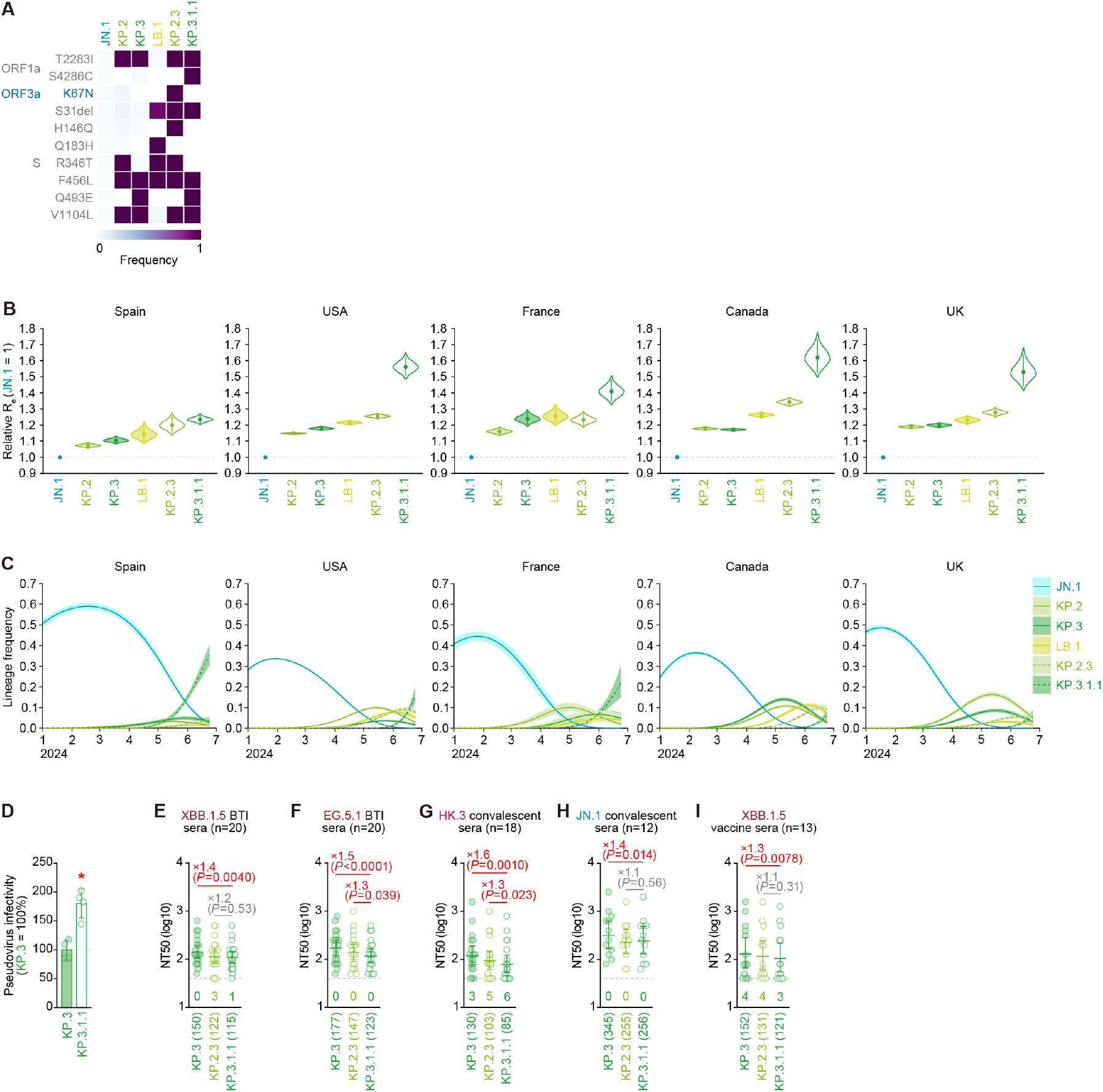
Virological features of KP.3.1.1. (**A**) Frequency of mutations in KP.3.1.1, and other lineages of interest. Only mutations with a frequency >0.5 in at least one but not all the representative lineages are shown. (**B**)Estimated relative R_e_ of the variants of interest in Spain, the USA, France, Canada, and the UK. The relative R_e_ of JN.1 is set to 1 (horizontal dashed line). Violin, posterior distribution; dot, posterior mean; line, 95% credible interval. (**C**)Estimated epidemic dynamics of the variants of interest in Spain, the USA, France, Canada, and the UK from January 1, 2024, to June 24, 2024. Countries are ordered according to the number of detected sequences of KP.3.1.1 from high to low. Line, posterior mean, ribbon, 95% credible interval. (**D**)Lentivirus-based pseudovirus assay. HOS-ACE2/TMPRSS2 cells were infected with pseudoviruses bearing each S protein of KP.3 and KP.3.1.1. The amount of input virus was normalized to the amount of HIV-1 p24 capsid protein. The percentage infectivity of KP.3.1.1 is compared to that of KP.3. The horizontal dash line indicates the mean value of the percentage infectivity of KP.3. Assays were performed in quadruplicate, and a representative result of four independent assays is shown. The presented data is expressed as the average ± SD. Each dot indicates the result of an individual replicate. Statistically significant differences versus KP.3 is determined by two-sided Student’s *t* tests and statistically significant difference (*P* < 0.05) versus KP.3 is indicated with red asterisk. (**E**-**I**) Neutralization assay. Assays were performed with pseudoviruses harboring the S proteins of KP.3, KP.2.3 and KP.3.1.1. The following convalescent sera were used: sera from fully vaccinated individuals who had been infected with XBB.1.5 (one 2-dose vaccinated donor, seven 3-dose vaccinated donors, six 4-dose vaccinated donors, five 5-dose vaccinated donors and one 6-dose vaccinated donor. 20 donors in total) (**E**); EG.5.1 (one 2-dose vaccinated donor, six 3-dose vaccinated donors, five 4-dose vaccinated donors, four 5-dose vaccinated donors and four 6-dose vaccinated donors. 20 donors in total) (**F**); individuals who had been infected with HK.3 (three 2-dose vaccinated donors, five 3-dose vaccinated donor, two 4-dose vaccinated donors, three 5-dose vaccinated donors, one 6-dose vaccinated donor and four donors with unknown vaccine history. 18 donors in total) (**G**) and individuals who had been infected with JN.1 (one 2-dose vaccinated donor, two 3-dose vaccinated donors, two 7-dose vaccinated donors and seven donors with unknown vaccine history. 12 donors in total) (**H**), and XBB.1.5 monovalent vaccination sera from fully vaccinated individuals who had been infected with XBB subvariants (after June, 2023) (13 donors) (**I**). Assays for each serum sample were performed in quadruplicate to determine the 50% neutralization titer (NT_50_). Each dot represents one NT_50_ value, and the geometric mean and 95% confidence interval are shown. The number in parenthesis indicates the geometric mean of NT_50_ values. The horizontal dash line indicates a detection limit (40-fold) and the number of serum donors with the NT_50_ values below the detection limit is shown in the figure (under the bars and dots of each variant). Statistically significant differences versus KP.3.1.1 were determined by two-sided Wilcoxon signed-rank tests, and p values are indicated in parentheses. The fold changes of NT50 versus KP.3.1.1 are indicated with “X”.

First, we estimated the relative effective reproduction number (R_e_) of KP.3.1.1 using a Bayesian multinomial logistic model^4^ based on genome surveillance data from Spain, the USA, France, Canada, and the UK, where this variant has spread as of June 2024 (**Figures 1B and 1C; Table S4**). In Spain, the R_e_ of KP.3.1.1 is over 1.2-fold higher than that of JN.1 and even higher than those of KP.2, KP.3, LB.1, and KP.2.3 (**Figure 1B**). Additionally, the other countries under investigation herein show higher R_e_ for KP.3.1.1. However, it must be noted there is the possibility of overestimation in these countries due to more limited KP.3.1.1 sequence numbers. These results suggest that KP.3.1.1 will spread worldwide along with other JN.1 sublineages.^2,3^

We then assessed the virological properties of KP.3.1.1 using pseudoviruses. The pseudovirus of KP.3.1.1 had significantly higher infectivity than that of KP.3 (**Figure 1D**). Neutralization of KP.3.1.1 was tested using i) convalescent sera after breakthrough infection (BTI) with XBB.1.5 or EG.5, ii) convalescent sera after the infection with HK.3 or JN.1, and iii) sera after monovalent XBB.1.5 vaccination. The 50% neutralization titer (NT_50_) against KP.3.1.1 was significantly lower than KP.3 (1.4–1.6-fold) in all four groups of convalescent sera tested (**Figures 1E–H**). KP.3.1.1 also showed a 1.3-fold lower NT_50_ against XBB.1.5 vaccine sera than KP.3 (**Figure 1I**).^1–3,5^ Moreover, KP.3.1.1 showed stronger resistance with a 1.3-fold lower NT_50_ with statistical significances to the convalescent sera infected with EG.5 and HK.3 than KP.2.3 (**Figures 1F and 1G**).

Altogether, KP.3.1.1 exhibited a higher R_e_, higher pseudovirus infectivity, and higher neutralization evasion than KP.3. These results align with our recent report that the JN.1 subvariants with S:S31del (e.g., KP.2.3 and LB.1) exhibited enhanced R_e_ and immune evasion compared to the other JN.1 subvariants without S:S31del (e.g., JN.1, KP.2, and KP.3),^3^ highlighting the evolutionary significance of S:S31del in the JN.1 lineages.

